# Pathogenicity of urinary tract infection *Escherichia coli* in *Caenorhabditis elegans*

**DOI:** 10.1101/858910

**Authors:** Masayuki Hashimoto, Yi-Fen Ma, Sin-Tian Wang, Chang-Shi Chen, Ching-Hao Teng

**Affiliations:** Institute of Molecular Medicine, College of Medicine, National Cheng Kung University, Tainan, Taiwan (ROC); Center of Infectious Disease and Signaling Research, National Cheng Kung University, Tainan, Taiwan (ROC); Institute of Basic Medical Sciences, College of Medicine, National Cheng Kung University, Tainan, Taiwan (ROC); Department of Biochemistry and Molecular Biology, College of Medicine, National Cheng Kung University, Tainan, Taiwan (ROC)

## Abstract

Uropathogenic *Escherichia coli* (UPEC) is a major bacterial pathogen that causes urinary tract infections (UTIs). Several virulence factors (VFs) in the bacteria have been associated with the pathogenicity. The mouse is an available UTI model for studying the pathogenicity; however, *Caenorhabditis elegans* represents as an alternative surrogate host for studying UPEC with the capacity for high-throughput analysis. Therefore, we established a simple assay for a UPEC infection model with *C. elegans* for large-scale screening. An *E. coli* culture to be tested and synchronized *C. elegans* were mixed in 96-well plates, and the pathogenicity was determined by comparison of the turbidity before and after incubation. A total of 133 clinically isolated *E. coli* strains, which included UTI-associated and fecal isolates, were applied to demonstrate the liquid pathogenicity assay. The *E. coli* isolates associated with UTIs showed higher pathogenicity in *C. elegans* than the fecal isolates, suggesting that the simple assay with *C. elegans* is useful as a UPEC infectious model. From the screening, VFs involved with iron acquisition (*chuA, fyuA*, and *irp2*) were significantly associated with high pathogenicity. *C. elegans* is a heme auxotroph, and iron homeostasis also serves innate immunity in *C. elegans*. We then evaluated whether the VFs in UPEC were involved in the pathogenicity. Mutants of *E. coli* UTI89 with defective iron acquisition systems were applied to a solid killing assay with *C. elegans*. As a result, the survival rate of *C. elegans* fed with the mutants significantly increased compared to when fed with the parent strain. To our knowledge, this is the first report of the involvement of iron acquisition in the pathogenicity of UPEC in a *C. elegans* model.

## Introduction

Urinary tract infections (UTIs) are very common [1], and the resulting direct medical cost is substantial [2]. Uropathogenic *E. coli* (UPEC) is one of the major etiologies of UTIs. A UPEC strain usually requires an array of virulence factors (VFs) with different functions to invade the urinary tract. VFs of pathogenic bacteria are potential antimicrobial targets and markers of the pathogen. However, different UPEC strains may harbor varying combinations of VFs, and most of the factors only exist in a fraction of UPEC strains. Thus, a combination of multiple VFs are required for developing effective and widely usable measures to prevent or treat UTIs caused by UPEC. Identifying how VFs contribute to UTIs and understanding their epidemiological distribution would facilitate the development of such novel strategies to manage the infection.

The mouse is generally used as an animal model for studying the pathogenicity of UPEC. From a technical, economical, and an ethical viewpoint, however, the model is not suitable for use in large-scale studies. The *Caenorhabditis elegans* model is cheap, easy to handle, and has been applied in various bacterial pathogenicity studies such as with *Pseudomonas aeruginosa* [3], *Enterococcus* [4], *Serratia marcescens* [5], and *E. coli* [6-12] including UPEC [9, 10]. For the UPEC study, the results revealed that the pathogenicity in *C. elegans* was associated with the number of VFs, which were identified as virulence genes involved in the pathogenicity in mammal [9]. Namely, the VFs involved in the infection in mammal is proposed to function in the *C. elegans* model, although the association had not been proven in a previous study using mutant VF.

In most of the aforementioned *C. elegans* models, synchronized *C. elegans* animals were incubated with a lawn of interested bacteria on a plate, and the survival rates of the animals were determined during the examination (solid killing assay) [3-7, 9-11]. The surviving *C. elegans* must be transferred to a fresh plate with the lawn every day during the pregnancy period to avoid affection of newborn *C. elegans* animals in the assessment. The copious handling for transferring of *C. elegans* makes it difficult to use in large-scale screening. Thus, a liquid killing assay was developed [8]. In this method, *C. elegans* incubated with the lawn of interested bacteria was harvested with liquid medium from the plate after incubation for 1 day, and the survivability of *C. elegans* in the liquid medium was observed in a time-dependent manner. To further simplify the assay, we recently developed another liquid pathogenicity assay to screen a transposon mutant library of *E. coli* o157 comprised of 17,802 mutants [12]. Pathogenicity was evaluated by measuring the OD_595nm_ value of a mixture of mutant *E. coli* and *C. elegans* in 96-well plates after 8 days of incubation.

In this study, we modified the liquid pathogenicity assay described above to apply it to 133 clinically isolated *E. coli*, including 83 UTI-associated isolates and 50 fecal isolates, and we analyzed the association between pathogenicity in *C. elegans* and the VFs in the isolates. In addition, we demonstrated that the iron acquisition systems associated with the pathogenicity of UPEC are involved in the virulence in *C. elegans*.

## Materials and methods

### Strain and culture of bacteria and nematodes

In this study, 81 *E. coli* isolates from patients with UTI-related symptoms from National Cheng Kung University Hospital (NCKUH) in Taiwan from January to April 2005 were studied as UTI-associated isolates [13]. A total of 49 *E. coli* fecal isolates from healthy volunteers were also collected from NCKUH between January and June 2006 [13]. In addition, *E. coli* UTI89 and CFT073 were used as model UPEC, and *E. coli* K-12 MG1655 as a model non-pathogenic *E. coli*. These *E. coli* stains were cultured in lysogeny broth (LB) at 37 °C with vigorous shaking [14].

*C. elegans* N2 as wild type was used for the solid killing assay. The worm was maintained on NGM agar plates using the standard laboratory *E. coli* strain OP50 as described previously [15]. *C. elegans glp-4*(*bn2*ts) was used for the liquid pathogenicity assay. The temperature-sensitive strain was maintained at 15 °C for animal preparation.

### Phylogenetic and virulence factor analyses

For the 133 *E. coli* used in this study, the phylogenetic group and 31 VFs identified previously in UPEC were determined by PCR as described previously [16-20]. The primers used in these analyses are listed in S1 Table.

### Mutant construction of *E. coli* UTI89

Gene deletion in *E. coli* UTI89 was performed by applying the lambda *red* system with pKD46 as described previously [21]. To delete *chuA*, the Cm^R^ cassette on pKD3 was PCR-amplified with primers (chuA-P1 and chuA-P2), and the parent strain with pKD46 was transformed with the PCR product to obtain *chuA* replaced with the Cm^R^ cassette. The deletion of *fyuA* was performed similarly with *chuA* replaced with the Km^R^ cassette amplified by PCR with pKD4 as template, and fyuA-P1 and fyuA-P2 as primers. A plasmid expressing Flp (pCP20) was used to remove the antibiotic markers flanked by FLPs. For *entA* deletion, the Δ*entA*::Km region from the KEIO collection (JW0588-KC) was amplified with 165-1 and 165-2, and applied for the lambda *red* recombination [22]. The genetic structures of all the recombinations were confirmed by PCR. *E. coli* UTI89 and its derivatives that were used for solid killing assay are listed in S2 Table.

### Liquid pathogenicity assay

The pathogenicity assay method has been previously described in detail [12]. In brief, *C. elegans glp-4*(*bn2*ts), which is temperature sensitive for pregnancy, was used to fix *C. elegans* generation during the assay. *E. coli* cells at an OD_600nm_ of 0.2 that were in the stationary phase cultured in LB medium were transferred to a 96-well plate. Next, the plate was centrifuged (3,700 rpm, 10 min, 25 °C), the resulting pellet of *E. coli* cells collected, and the supernatant discarded. Synchronized *C. elegans* in late L4 stage were prepared as described previously [23], and approximately 30 animals in 200 µl of S medium were added to the *E. coli* pellet. The bacteria and animals were co-incubated at 25 °C with shaking (100 rpm) for 8 d, at which time the OD_592nm_ was measured and normalized with the value at 0 d. The experiments were performed independently three times.

### Solid killing assay

The survivability of the *C. elegans* N2 strain feeding the *E. coli* UTI89 wild type or its mutants was measured as described previously [7], but with some modification. Briefly, *E. coli* strains were cultured in LB medium at 37 °C until at an OD_600nm_ of 2.0, and 30 µl of the culture was spread on a 5.0 cm NGM agar plate and incubated at 37 °C overnight. On the next day, approximately 50 synchronized *C. elegans* N2 animals in late L4 stage were inoculated on each plate and incubated at 20 °C for 10 d. During the assay, the survivability of the *C. elegans* on the plate was measured daily, and living animals were transferred to a fresh plate. Nematoda that did not respond to gentle prodding were scored as dead. Animals that crawled off the plate were censored. The experiments were performed with approximately 100 worms total per *E. coli* strain.

### Statistical analysis

The statistical analysis of the phylogenetic and virulence factor data was performed with Fischer’s exact test. The Mann–Whitney U test was used for analysis of the liquid pathogenicity data. For the solid killing assay, the Mantel–Cox log-rank test was used to assess the statistical significance of the difference in survival of the *C. elegans*. These analyses were done by using GraphPad Prism (GraphPad Software) version 7.0.

## Results

### Epidemiological characterization of the clinically isolated *E. coli* strains

The phylogenetic group and VFs of the strains used in this study were determined to characterize the strains. Eighty-three *E. coli* strains isolated from UTIs including the prototype UPEC strains UTI89 and CFT073, and 50 *E. coli* fecal isolates including *E. coli* MG1655 as commensal strains were used. Based on the syndromes of these UTI-associated strains, they were further divided into the lower UTI (cystitis associated strains), upper UTI (pyelonephritis associated strains), and urosepsis groups (S3 Table). The phylogenetic groups of the clinical isolates used in this study are indicated in Table 1. Most of the UTI-associated isolates belonged to the B2 group, which mainly consists of extraintestinal pathogenic strains that display a high concentration of VFs [24]. In contrast, most of the fecal isolates are associated with phylogenetic group A, whose member strains are often devoid of extraintestinal VFs [25]. The distribution of 31 genes identified as uropathogenic VFs [17-20] were determined in the 133 *E. coli* isolates (Table 1). Eighteen genes showed significantly higher distribution in the UTI-associated isolates than was observed in the fecal isolates, while the level of *ibeA* was higher in the fecal isolates. The total number of VFs identified was significantly higher in the UTI-associated isolates than in the fecal isolates (Fig. 1).

**Table 1.**
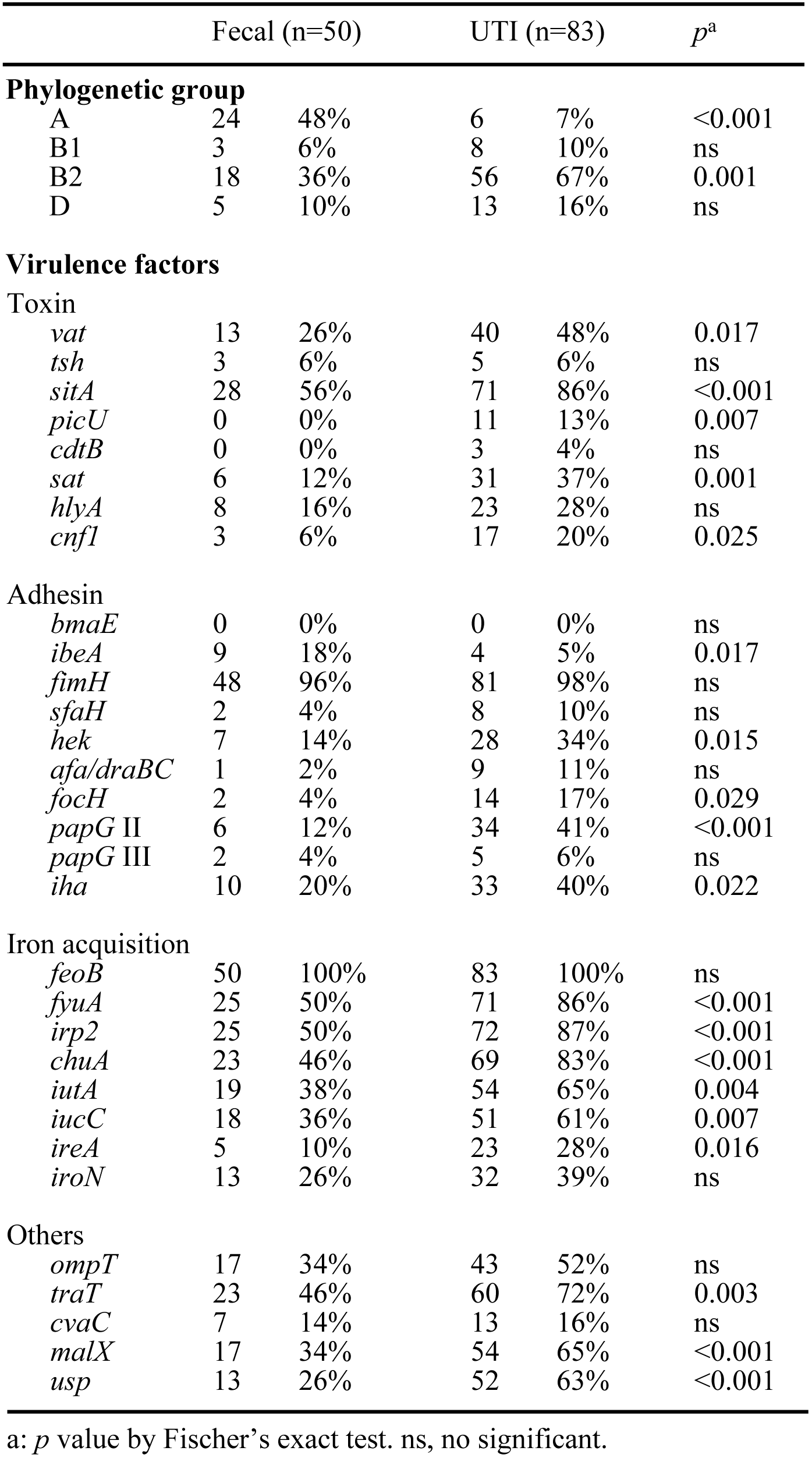
Characterization of the 133 *E. coli* clinical isolates.

**Fig 1.**
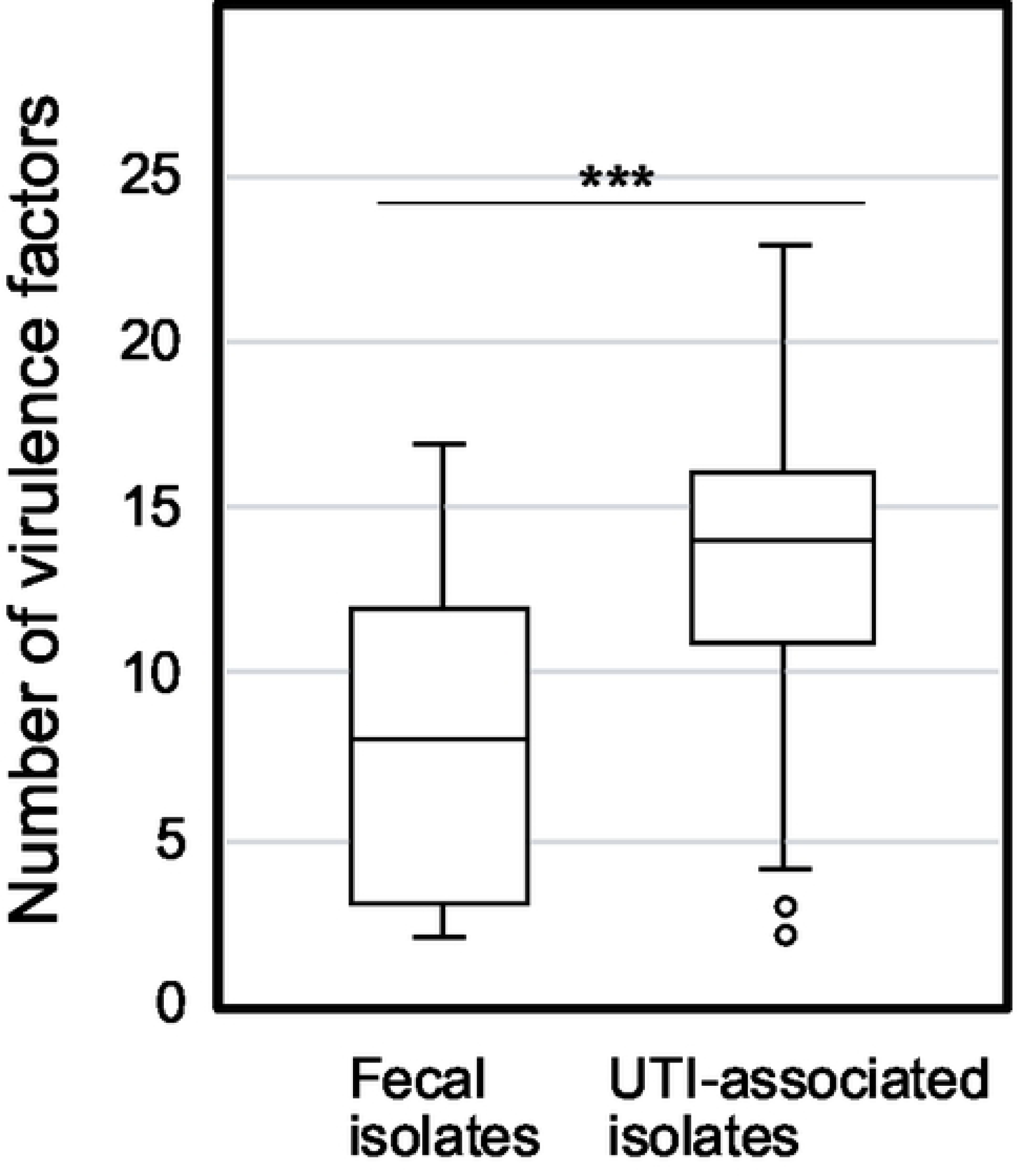
Number of virulence factors in clinically isolated *E. coli*. The boxplot indicates the number of VFs in fecal and UTI-associated isolates. The numbers of VFs between fecal and UTI-associated isolates (average 8.1 and 13.2, respectively) show significant difference by the Mann–Whitney U test (***, *p*<0.001).

### Liquid pathogenicity assay of *E. coli* with *C. elegans*

The pathogenicity of the *E. coli* isolates was determined by a liquid assay with *C. elegans* in which both species were incubated in S medium on 96-well plate for 8 d, followed by OD_595nm_ measurement to determine *E. coli* survival. When a tested *E. coli* strain is non-pathogenic, the turbidity will be low after incubation because the *E. coli* cells are fed on by *C. elegans*. In contrast, when a *C. elegans* is not healthy with a pathogenic *E. coli*, the bacterial cells survive and show higher turbidity. As shown in Fig 2, the pathogenicity of UTI-associated isolates was significantly higher than that of the fecal isolates. The result is consistent with the higher number of VFs in UTI-associated isolates (Fig 1) and suggests that the *C. elegans* model reflects the pathogenicity in patients. In comparing symptoms, the isolates associated with lower and upper UTIs showed significantly higher pathogenicity than the fecal isolates, while the pathogenicity of urosepsis was comparable with that of commensal *E. coli* (S1 Fig). Since the number of VFs in urosepsis was significantly higher than that in the fecal isolates (S2 Fig), the VFs for urosepsis might not be involved in the pathogenicity in *C. elegans*. Regarding the phylogenetic groups, the pathogenicity of groups B1, B2, and D were significantly higher than that of group A (S1 Fig), although the number of VFs in group B1 was comparable with group A (S2 Fig). Perhaps, isolates in group B1 bear unidentified VFs for *C. elegans*. The association between each VF and pathogenicity was calculated in Table 2. Seven VFs showed a significant association with the pathogenicity in *C. elegans*, and three of the seven VFs belonged to the iron acquisition group while the other four were in the other-group. These results suggest that the VFs for iron acquisition participate in pathogenicity in *C. elegans*.

**Table 2.**
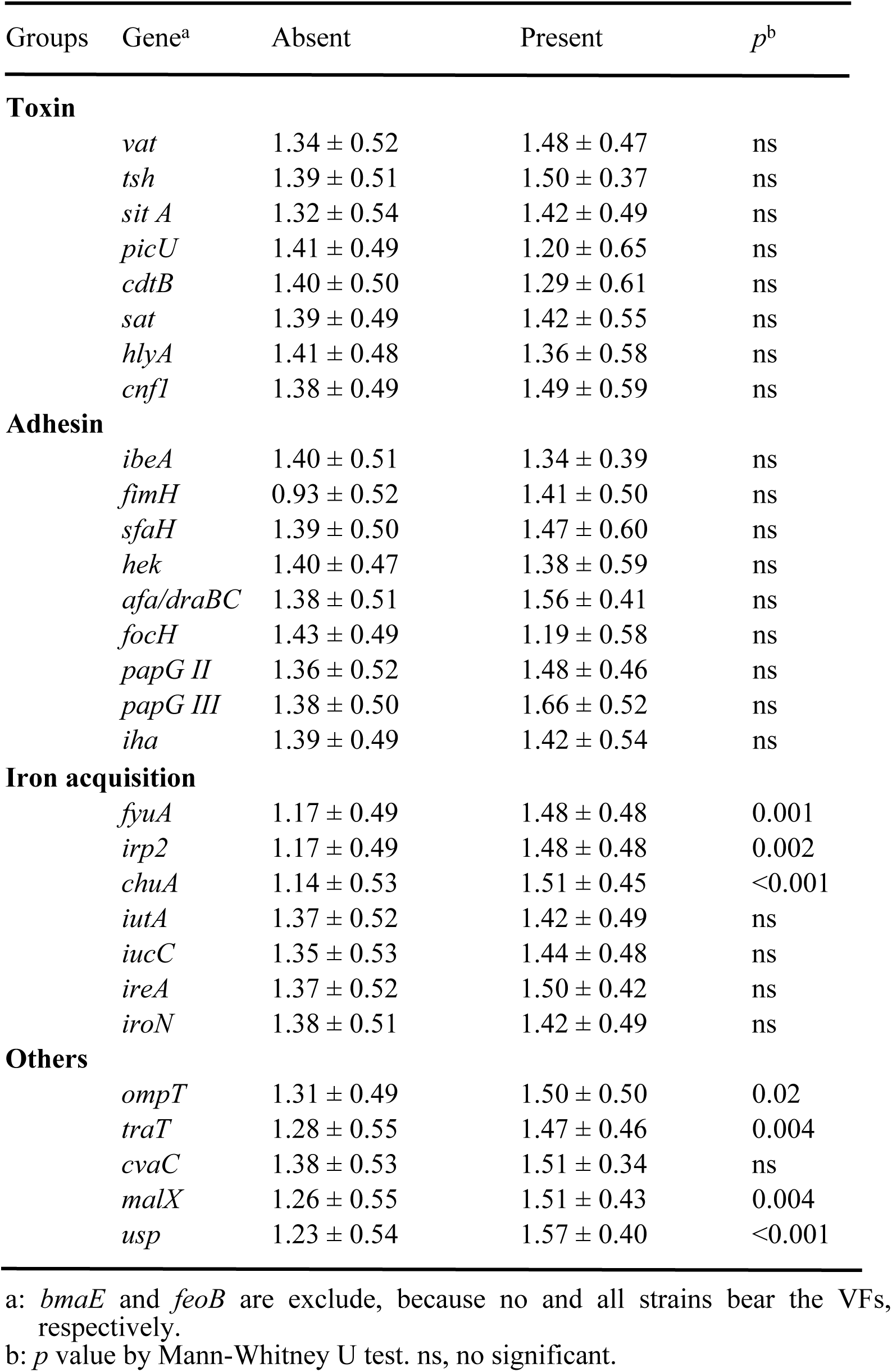
The association between pathogenicity and virulence factor.

| Groups | Gene<sup>a</sup> | Absent | Present | $p^b$ |
| --- | --- | --- | --- | --- |
| **Toxin** |  |  |  |  |
|  | *vat* | 1.34 ± 0.52 | 1.48 ± 0.47 | ns |
|  | *tsh* | 1.39 ± 0.51 | 1.50 ± 0.37 | ns |
|  | *sit A* | 1.32 ± 0.54 | 1.42 ± 0.49 | ns |
|  | *picU* | 1.41 ± 0.49 | 1.20 ± 0.65 | ns |
|  | *cdtB* | 1.40 ± 0.50 | 1.29 ± 0.61 | ns |
|  | *sat* | 1.39 ± 0.49 | 1.42 ± 0.55 | ns |
|  | *hlyA* | 1.41 ± 0.48 | 1.36 ± 0.58 | ns |
|  | *cnfI* | 1.38 ± 0.49 | 1.49 ± 0.59 | ns |
| **Adhesin** |  |  |  |  |
|  | *ibeA* | 1.40 ± 0.51 | 1.34 ± 0.39 | ns |
|  | *fimH* | 0.93 ± 0.52 | 1.41 ± 0.50 | ns |
|  | *sfaH* | 1.39 ± 0.50 | 1.47 ± 0.60 | ns |
|  | *hek* | 1.40 ± 0.47 | 1.38 ± 0.59 | ns |
|  | *afa/draBC* | 1.38 ± 0.51 | 1.56 ± 0.41 | ns |
|  | *focH* | 1.43 ± 0.49 | 1.19 ± 0.58 | ns |
|  | *papG II* | 1.36 ± 0.52 | 1.48 ± 0.46 | ns |
|  | *papG III* | 1.38 ± 0.50 | 1.66 ± 0.52 | ns |
|  | *iha* | 1.39 ± 0.49 | 1.42 ± 0.54 | ns |

Iron acquisition
|  |  |  |  |
| --- | --- | --- | --- |
| *fyuA* | 1.17 ± 0.49 | 1.48 ± 0.48 | 0.001 |
| *irp2* | 1.17 ± 0.49 | 1.48 ± 0.48 | 0.002 |
| *chuA* | 1.14 ± 0.53 | 1.51 ± 0.45 | <0.001 |
| *iutA* | 1.37 ± 0.52 | 1.42 ± 0.49 | ns |
| *iucC* | 1.35 ± 0.53 | 1.44 ± 0.48 | ns |
| *ireA* | 1.37 ± 0.52 | 1.50 ± 0.42 | ns |
| *iroN* | 1.38 ± 0.51 | 1.42 ± 0.49 | ns |

Others
|  |  |  |  |
| --- | --- | --- | --- |
| *ompT* | 1.31 ± 0.49 | 1.50 ± 0.50 | 0.02 |
| *traT* | 1.28 ± 0.55 | 1.47 ± 0.46 | 0.004 |
| *cvaC* | 1.38 ± 0.53 | 1.51 ± 0.34 | ns |
| *malX* | 1.26 ± 0.55 | 1.51 ± 0.43 | 0.004 |
| *usp* | 1.23 ± 0.54 | 1.57 ± 0.40 | <0.001 |

a: *bmaE* and *feoB* are exclude, because no and all strains bear the VFs, respectively.
b: *p* value by Mann-Whitney U test. ns, no significant.

**Fig 2.**
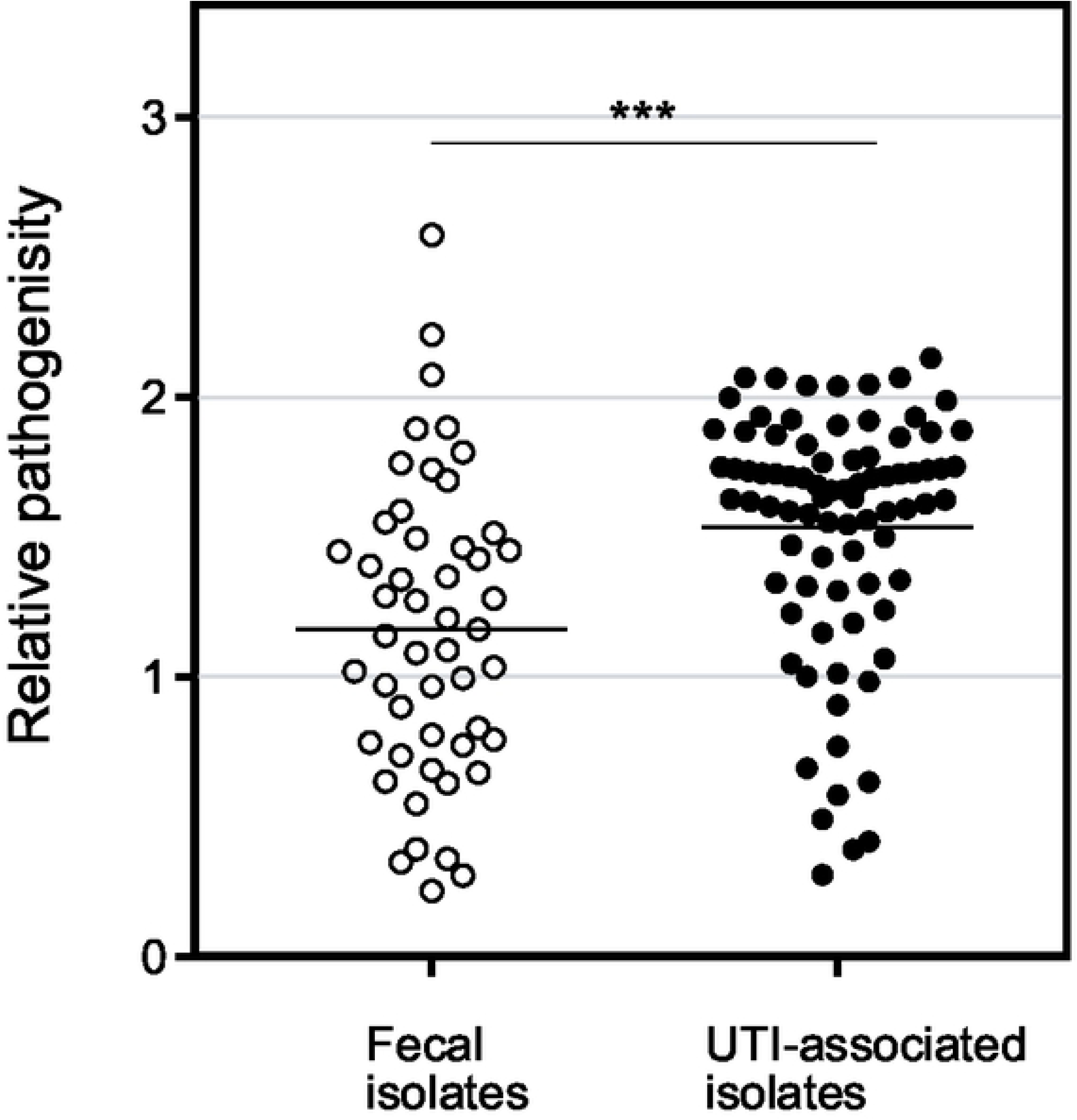
Pathogenicity of *E. coli* clinical isolates in liquid killing assay with *C. elegans.* The pathogenicity (relative OD_592nm_ value in liquid assay) of the fecal and UTI-associated isolates were plotted. The horizontal bars represented the mean values. The results were analyzed statically by the Mann–Whitney U test. (**, *p*<0.01). The pathogenicity of UTI-associated isolates was significantly higher than that of fecal isolates.

### Iron acquisition of UPEC involved in the pathogenicity in *C. elegans*

To investigate whether VFs for iron acquisition (*chuA, irp2, fyuA*) are involved in the pathogenicity in *C. elegans*, UPEC mutants defective in the genes were applied to the animal model. One of the VFs, *chuA*, is a receptor for heme acquisition [26] and has been reported to be involved in kidney infection in the mouse model [27]. The other two genes are for yersiniabactin as a siderophore, and *iro2* is for biosynthesis and *fyuA* is a receptor for the siderophore [28, 29]. The genomic sequences of the UPEC strains used in this study, UTI89 and CFT073, have been published [30, 31]. Although both of the strains carry the three VFs, only UTI89 was used for further study because it produces yersiniabactin while CFT073 does not [32]. Specifically, UTI89 uses heme and produces the three siderophores yersiniabactin, enterobactin, and salmochelin [32]. As described above, the genes for yersiniabactin show significant association with the pathogenicity. On the other hand, the genes for enterobactin were not analyzed in the liquid pathogenicity assay because it is found in most *E. coli* strains. However, because the involvement of enterobactin in mammal infection has been reported in UPEC [33], enterobactin was also investigated. In addition, the mutant defective in enterobactin synthesis (Δ*entA*) does not produce either enterobactin and salmochelin, because salmochelin is derived from enterobactin [34]. Multiple iron acquisition systems are generally identified in UPEC, and the redundancy is important for infection [35]. Therefore, the mutants combinatorially defective in heme acquisition (Δ*chuA*), yersiniabactin acquisition (Δ*fyuA*), and enterobactin and salmochelin production (Δ*entA*) were constructed and applied in the solid killing assay with *C. elegans*. As a result, all the mutant strains defective in the VFs for iron acquisition showed lower pathogenicity than wild type (Fig 3). The pathogenicity of a single mutation of *chuA* and *fyuA*, which were associated with pathogenicity in the liquid assay, were attenuated. A double mutant (Δ*chuA* Δ*fyuA*) showed lower pathogenicity than each of the single mutants. The pathogenicity of the *entA* single mutation was also decreased compared to wild type. Furthermore, although the difference in the pathogenicity of triple mutant Δ*chuA ΔfyuA ΔentA* was not statically significant when compared with the double mutant, the finding still supported the trend in decreasing pathogenicity. Taken together, these results demonstrated that iron acquisition is involved in the pathogenicity of UPEC in *C. elegans*. However, another mechanisms might also be involved, because the pathogenicity of the triple mutant is much higher than that of OP50, which generally shows over 50% survival for two weeks.

**Fig 3.**
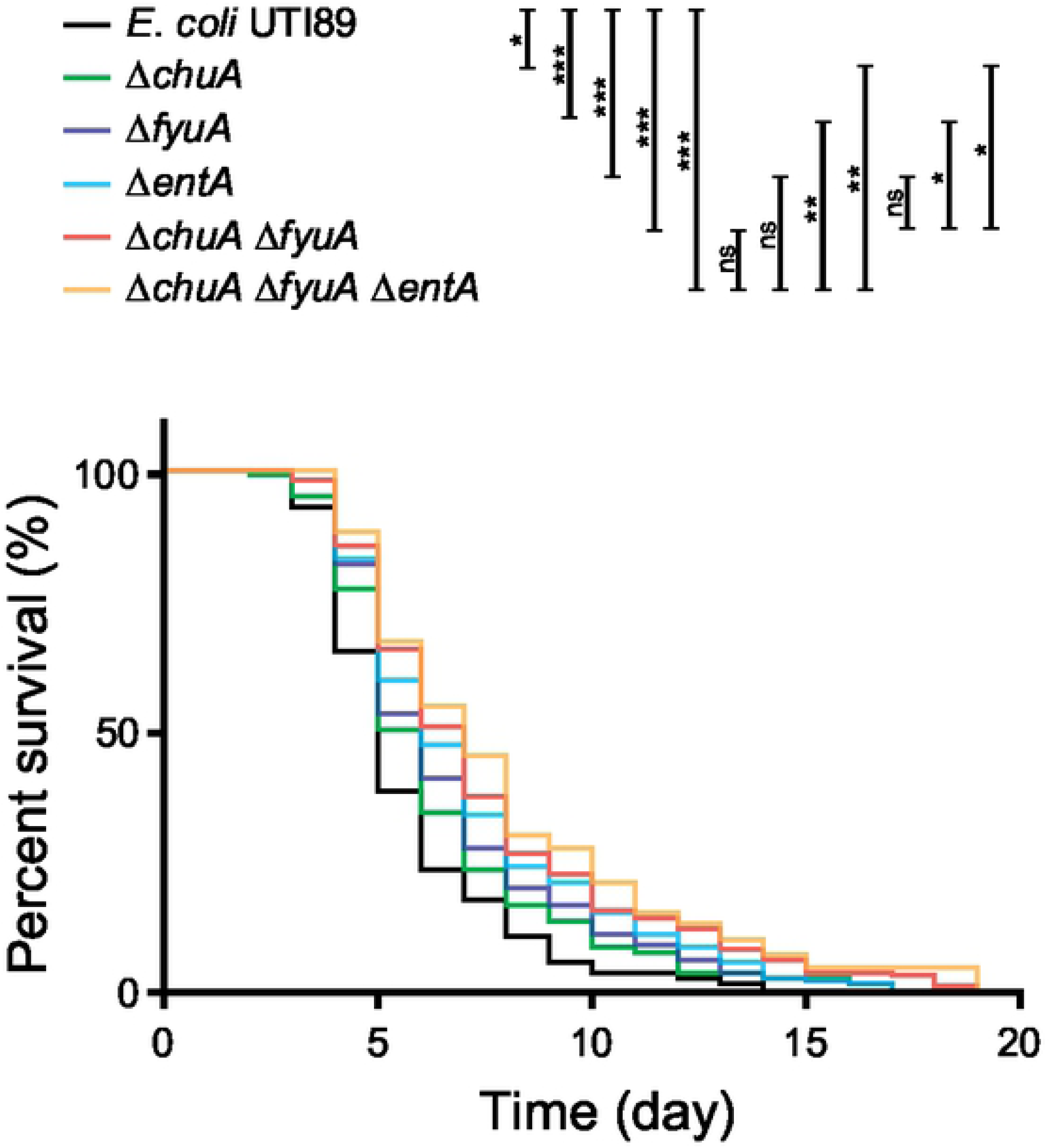
UPEC mutants defective in iron acquisition decreased the pathogenicity in *C. elegans* in the solid killing assay. The survivability of *C. elegans* (*n*>100) feeding on the wild type and mutants of *E. coli* UTI89 defective in iron acquisition, which shows association with pathogenicity in the liquid pathogenicity assay. The results were analyzed statically by the Mantel–Cox log-rank test (*, *p*>0.05; **, *p*>0.01; ***, *p*>0.001; ns, not significant).

## Discussion

### Liquid pathogenicity assay

The pathogenicity of *E. coli* associated with UTIs was determined using *C. elegans*. Although several reports have been published on bacterial pathogenicity using *C. elegans*, the studies used the *C. elegans* killing assay, which is based on counting the animals to determine survivability on a solid or in a liquid medium [3-7, 9-11]. Here, we adopted a liquid pathogenicity assay in which *C. elegans* and *E. coli* were co-incubated, and the turbidity was compared before and after the incubation. Although the method has been applied for a mutant library derived from one parent strain [12], this is the first report to apply it to numerous clinical isolates. Since the clinically isolated *E. coli* associated with UTIs showed significantly higher pathogenicity than the commensal strains, the results of the simple method in this study reflected the pathogenicity in the patient (Fig 2). In addition, we found that the numbers of VFs belonging to toxin, adhesin, and iron acquisition were detected with higher frequency in UTI-associated isolates than in feces isolates (Table 1). However, only 7 VFs belonging to the iron acquisition and other-group showed significant association with high pathogenicity in *C. elegans* (Table 2). Perhaps the VFs toxin and adhesin are not involved in the pathogenicity in *C. elegans*. Alternatively, the sensitivity of the method used in this study may be not enough to detect the difference.

### Iron acquisition

To demonstrate whether the iron acquisition systems participate in the pathogenicity of UPEC in *C. elegans, E. coli* UTI89 mutants defective in the VFs were constructed and applied in the *C. elegans* solid killing assay. The statistical associations between the VFs and pathogenicity in *C. elegans* have been reported in UPEC [6, 9, 10]. However, it has not been reported that iron acquisition is involved in pathogenicity in the *C. elegans–E. coli* model, while it is important in infection in mammal. To the best of our knowledge, this is the first study to demonstrate that iron acquisition in *E. coli* is involved in the pathogenicity in *C. elegans*.

Iron is an essential element for all organisms. Iron homeostasis in mammal serves as an innate immune response to prevent bacterial infection. Therefore, iron acquisition in bacteria is important for virulence [36]. In *C. elegans*, many ortholog genes for iron homeostasis in mammal were identified, and the mammalian iron-metabolism is generally conserved [37]. In this context, iron deprivation prolonged the survivability of *C. elegans* infected with *Salmonella enterica* serovar Typhimurium [38]. The expression of *smf-3* and *smf-1* in *C. elegans* which are orthologs of divalent-metal transporter 1 (DMT-1) in mammal were induced by exposure to *Staphylococcus aureus*, and mutants of the transporters showed hypersensitivity to the pathogen [39]. In addition, a ferritin homolog *ftn-2* involved in cellular iron storage has been shown to be necessary for the full protective response to *E. coli* and *S. aureus* [40]. Consequently, iron homeostasis in *C. elegans* also serves as innate immunity.

In this study, the association between seven iron acquisition genes and pathogenicity in *C. elegans* were analyzed (Table 2), and *chuA* for heme acquisition, *irp2* for yersiniabactin synthesis, and *fyuA* for its receptor were found to be involved in the pathogenicity (Fig 3). Heme is the most abundant iron source in mammal, and it is an important iron source for bacterial infection. Two heme acquisition systems in UPEC (*chuA* and *hma*) were identified as VFs for mammal infection [35]. The expression of *chuA* was upregulated during the infection while that of *hma* was not [41, 42]. It appears that *chuA* plays an important role in UTIs. Since the worm is heme auxotroph [43], it is consistent with that heme acquisition by the pathogen affects the infection. In addition, while four siderophores have been identified in UPEC, UTI89 which was used in this study produces enterobactin, salmochelin, and yersiniabactin, but not aerobactin [32]. Enterobactin is a major iron acquisition system that is produced by most *E. coli*; however, it is sequestrated by lipocalin 2 during mammal infection [44]. On the other hand, salmochelin is a glucosylated enterobactin that is not bound by lipocalin 2, and can therefore escape the sequestration [45]. The involvement of enterobactin in *C. elegans* infection was also investigated in this study, because the worm does not produce lipocalin 2 to sequestrate enterobactin. Namely, UTI89 Δ*entA* Δ*fyuA*, which does not produce enterobactin and salmochelin nor uptake yersiniabactin, is defective in any siderophores. Available iron is limited in the urinary tract, and iron acquisition by bacteria is essential for the infection. Furthermore, multiple iron acquisition systems showed different roles in infection, so the redundancy is important for successful infection [35]. In this study, the combination of mutations for multiple iron acquisition systems resulted in decreasing pathogenicity in *C. elegans*, which is consistent with the importance of redundancy in mammal infection. In addition to the iron acquisition by siderophore, it has been recently reported that copper acquisition by yersiniabactin is also involved in UTIs [46]. While further study is necessary to investigate the molecular mechanism of iron acquisition in UPEC infection in *C. elegans*, availability of *C. elegans* for the model is demonstrated here.

## Acknowledgments

We are grateful to WP Zhang for wonderful technical assistance.

## Supplementary information

**S1 Fig. The pathogenicity of *E. coli* clinical isolates in different symptoms and phylogenetic groups.** The pathogenicity (relative OD_592nm_ value in liquid assay) of the 133 stains were plotted in different symptoms (panel A) and phylogenetic groups (panel B). The horizontal bars represented the mean values. The results were analyzed statically by Mann-Whitney U test (**, *p*<0.01; ***, *p*<0.001; ns, not significant).

**S2 Fig. Number of virulence factors in clinically isolated *E. coli*.** The box-plots indicate the number of VFs in each symptom (panel A) and in each phylogenetic group (panel B). The data was statically analyzed by Mann-Whitney U test (***, *p*<0.001; ns, not significant different).

**S1 Table. Primers used in this study.**

**S2 Table. *E. coli* strains used for solid killing assay.**

**S3 Table. Symptoms of clinically isolated 133 *E. coli* strains.**

